# Starch can expedite the screening for bacterial aflatoxin degraders

**DOI:** 10.1101/2023.09.27.559811

**Authors:** Natalie Sandlin, Babak Momeni

## Abstract

Aflatoxins (AFs) are secondary fungal metabolites that contaminate common food crops and are harmful to humans and animals. The ability to degrade or remove aflatoxins from common feed commodities will improve health standards and counter the economic drain inflicted by AF contamination. Bioremediation is a promising solution to AF contamination because of its low cost and few undesired environmental side-effects. Identifying new degrader species is highly beneficial in that it can offer alternatives to overcome the limitations of existing biodegraders, such as narrow working conditions and low degradation rates. Here, we screen several environmental isolates for their AF detoxification ability, using aflatoxin G_2_. We use different carbon sources (glucose and starch) as isolation and culturing media to examine the effect of the environment on degradation ability. Strains isolated in media with starch as the primary carbon source showed a higher percentage of good AF degraders, 16% compared to 2% when glucose was the primary carbon source. Additionally, the majority of species isolated in glucose medium exhibited improved degradation efficiency when moved into starch medium, with one isolate improving degradation levels from 30% to 70%. Our starch screen also revealed three previously unidentified AF degrader bacterial species. Good aflatoxin G_2_ degraders also appear to perform well against aflatoxin B_1_. Overall, for AF degradation, starch medium expedites the screening process and generally improves the performance of isolates. We thus propose that using starch as the carbon source is a promising means to identify new AF degraders in the environment.

## Introduction

Aflatoxins (AFs) are secondary fungal metabolites produces by *Aspergillus* spp. that ubiquitously contaminate common food and feed crops (1, 2). Consumption of contaminated foods by animals and humans pose serious complications to health including immunodepression, cancers, and hormone imbalances (3). Current physical and chemical methods for decontamination suffer from high costs, low reproducibility and efficiency, and the potential to diminish nutrients in the food (4–7). A more promising solution for AF decontamination is through biological methods, e.g. bioremediation using microbes (8–11). Bioremediation has the potential to offer lower cost, increased efficiency, and safer application in agriculture.

A number of bacterial and fungal species have been identified as AF degraders (10, 12–14), yet none have been effectively implemented for commercial use. The issues currently faced in using these species is in their narrow working conditions and unknown degradation mechanisms (15, 16). Therefore, it is highly beneficial to identify new degraders and their mechanisms of degradation to find species with broader working conditions and possibly higher and faster degradation rates. The process of finding new degrader species has previously been achieved through large screens testing environmental isolates either on the target toxin itself or compounds with a similar structure, such as coumarin in place of AF (12). However, such screens can be costly and result in limited numbers of degraders identified.

Here, we sought to implement a new screen that uses the ability of a microbe to grow on a more complex carbon source, starch, as an indication of its potential to degrade aflatoxin, also a complex carbon. Growth on starch is a good choice because starch is safe to handle, cost-effective, and allows rapid screening of isolates. Previous work in our lab has shown that simply changing microbes from a medium with glucose to a medium with starch as its sole carbon source increased degradation performance by strains (17). The presence of starch in the medium, compared to other carbon sources such as glucose, has also been shown to improve the AF degradation performance by fungus *Aspergillus niger* (18) and bacterium *Myroides odoratimimus* (19). Building on this background, we first screened environmental bacterial isolates from several sources for their ability to grow on starch defined medium and then tested for aflatoxin degradation by those able to grow in this medium. In parallel, we screened isolates on glucose defined medium prior to testing for degradation to determine differences in the strains obtained from different carbon sources. We used the native fluorescence of aflatoxin G_2_ (AFG_2_) to quantitatively characterize aflatoxin degradation by isolates. This degradation assay showed that a higher percentage of strains isolated on starch had degradation capability compared to those isolated on glucose. Additionally, the degradation performance of glucose isolates improved when tested in a starch environment in the degradation assay. These results indicate that starch can be utilized as a cost-effective screening tool for aflatoxin degraders and that environmental conditions such as carbon source can play a significant role in degradation rates.

## Results

### Selection on starch identified a greater percentage of good aflatoxin degraders

Environmental samples were taken from various locations in the surrounding area to broadly screen different species and environments for natural AF degraders: soil, leaf, sidewalk, doorknobs, and phone screens. After initial culturing on a rich solid (agar) medium, individual colonies (distinguished by different colony morphologies) were inoculated in a defined medium containing either starch or glucose as the sole carbon source (Fig. 1A). Out of 50 isolates tested for growth in starch, 26 were culturable and we subsequently tested them for aflatoxin degradation via our fluorescent AF degradation assay (Table 1). Only one starch isolate was unable to degrade AFG_2_ after 72 hours of testing and the 24 of the other 25 isolates showed degradation, with 4 having degradation efficiency >50% (Fig. 1B). Among glucose isolates, a larger percentage of tested isolates grew in the glucose medium (56 of 67), however, eight were unable to degrade AFG_2_ and only 1 showed degradation >50% (Table 1 and Fig. 1B), indicating that fewer active degraders arise in the glucose screen. The difference in good degraders that arose from the two screens is significantly favorable for the starch screen based on Fisher’s exact test, p=0.031 (Fig. 1B).

**Table 1.** Starch and glucose screen results.

| Selection medium | # of isolates | Grow in defined medium | Degradation performance |  |  |
| --- | --- | --- | --- | --- | --- |
|  |  |  | None | Poor (<50%) | Good (>50%) |
| Starch | 50 | 25 | 1 | 20 | 4 |
| Glucose | 67 | 55 | 8 | 46 | 1 |

**Figure 1.**
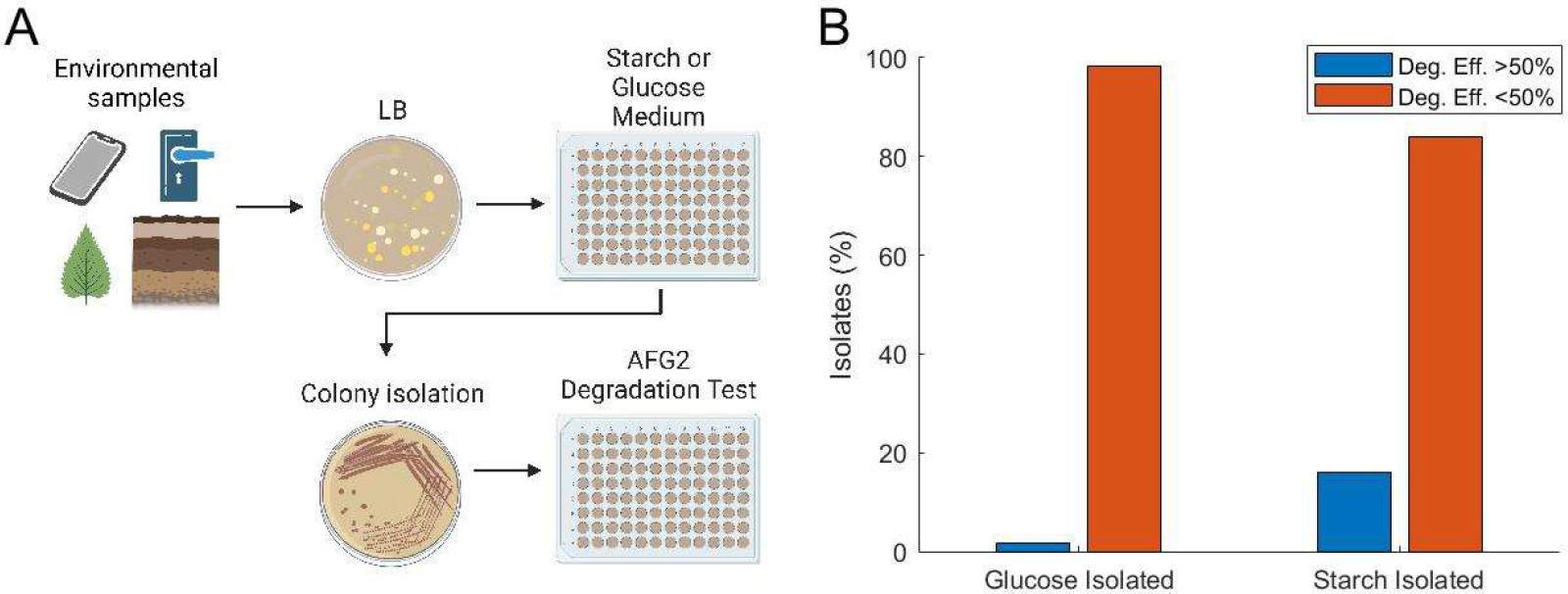
Profile of isolates from starch and glucose screens exhibits superior AFG2 degradation performance for those selected in media containing starch as the main carbon source. A) Experimental design schematic showing workflow from sample collection to isolate testing for AF degradation (Created with BioRender.com). B) Breakdown of degradation performance of environmental isolates based on selection medium. Blue bars show isolates that were able to degrade greater than 50% AF in the degradation assay and orange bars show isolates with less than 50% AF degradation. p = 0.031, Fisher’s exact test, comparing the number of good degraders arising from total isolates able to grow in glucose versus starch medium.

**Figure 2.**
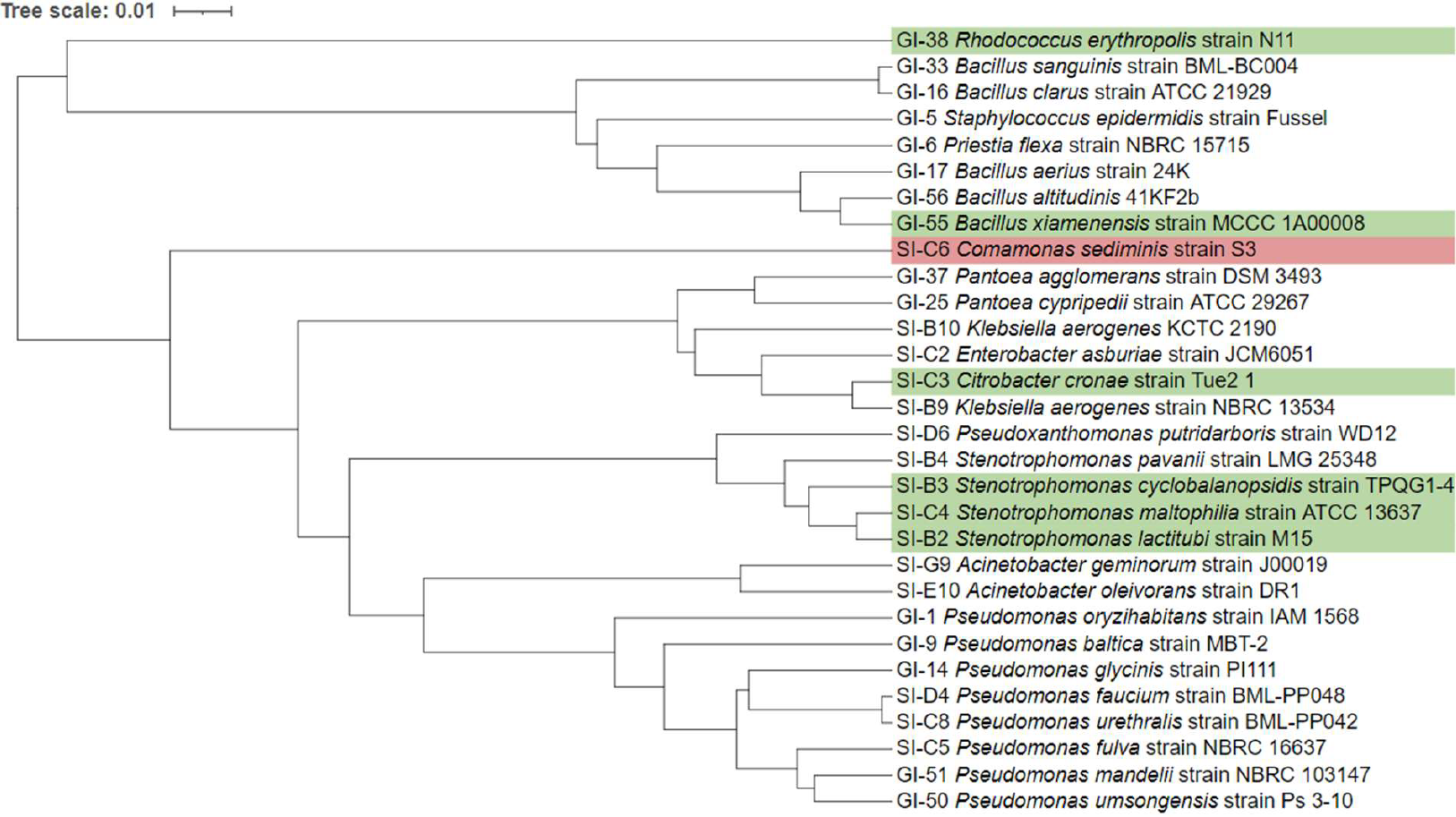
Good degraders are not phylogenetically distinct from poor degraders. Phylogenetic tree of glucose and starch isolates retrieved from GenBank database based on 16S rRNA sequences is shown. Green highlighted isolates are considered good degraders in starch medium, while red highlighted isolates are non-degraders in starch medium. GI and SI prefixes indicate those isolated in glucose and starch media, respectively.

### Newly identified aflatoxin degrading species arise from the starch screen

After testing for AF degradation by isolates in their isolation medium, 15 isolates from each screen were semi-randomly selected for further analysis to understand the general trends of the species that arose from each screen. Selected strains were chosen to represent the spectrum of degradation profiles, with representatives of the best, worst, and average performers. These selected isolates had their DNA extracted and PCR amplified for 16S rRNA sequencing to determine strain identity (Table 2). Of the glucose isolates analyzed, species of *Pseudomonas, Staphylococcus, Bacillus*, and *Pantoea* were present, all genera having previously been identified as aflatoxin degraders (20–27). Of starch isolates, while previously identified aflatoxin degrader genera were present, we also found species of *Citrobacter* and *Acinetobacter* which have not been previously implicated as aflatoxin degraders. To our knowledge, this is the first report of species in these two genera to possess aflatoxin degradation ability, although species from these two genera have been shown to degrade other mycotoxins: an *Acinetobacter* sp. isolated from soil degraded zearalenone (28), and a *Citrobacter* sp. isolated from soil degraded deoxynivalenol (29). Some of the identified isolates match previous literature at the genus level, and based on 16S rRNA sequencing we are not able to confirm or rule out if these isolates match the previously identified AF-degrading species. Overall, the starch screen resulted in newly identified degrader species while the glucose screen did not.

**Table 2.** Isolate identification and degradation profile. Isolates were identified through 16S rRNA sequencing. Identification shown in the table are closest match through BLASTn. Order of strains is based on performance in starch medium.

|  | Isolate | 16S rRNA identification | Isolated from | Live degradation (%) |  |
| --- | --- | --- | --- | --- | --- |
|  |  |  |  | In starch | In glucose |
| Starch Isolates | SI-C4 | *Stenotrophomonas maltophilia* | Soil | 64 | 13 |
|  | SI-B3 | *Stenotrophomonas cyclobalanopsidis* | Soil | 63 | 28 |
|  | SI-C3 | *Citrobacter cronae* | Soil | 59 | 11 |
|  | SI-B2 | *Stenotrophomonas lactitubi* | Soil | 53 | 39 |
|  | SI-E10 | *Acinetobacter oleivorans* | Soil | 31 | 13 |
|  | SI-C2 | *Enterobacter asburiae* | Soil | 29 | 27 |
|  | SI-C5 | *Pseudomonas fulva* | Soil | 22 | 29 |
|  | SI-D4 | *Pseudomonas faucium* | Tree trunk | 20 | 8.1 |
|  | SI-B10 | *Klebsiella aerogenes* | Soil | 20 | 17 |
|  | SI-B9 | *Klebsiella aerogenes* | Soil | 20 | 17 |
|  | SI-G9 | *Acinetobacter geminorum* | Sidewalk | 18 | 13 |
|  | SI-D6 | *Pseudoxanthomonas putridarboris* | Doorknob | 16 | 31 |
|  | SI-C8 | *Pseudomonas urethralis* | Soil | 11 | 11 |
|  | SI-B4 | *Stenotrophomonas pavanii* | Soil | 10 | 25 |
|  | SI-C6 | *Comamonas sediminis* | Soil | 3.8 | 19 |
| Glucose Isolates | GI-38 | *Rhodococcus erythropolis* | Soil | 84 | 82 |
|  | GI-55 | *Bacillus xiamenensis* | Leaf | 72 | 30 |
|  | GI-1 | *Pseudomonas oryzihabitans* | Phone screen | 47 | 34 |
|  | GI-5 | *Staphylococcus epidermidis* | Phone screen | 40 | 0.0 |
|  | GI-6 | *Priestia flexa* | Doorknob | 39 | 35 |
|  | GI-9 | *Pseudomonas baltica* | Leaf | 39 | 33 |
|  | GI-56 | *Bacillus altitudinis* | Soil | 38 | 35 |
|  | GI-50 | *Pseudomonas umsongensis* | Snow | 35 | 32 |
|  | GI-33 | *Bacillus sanguinis* | Soil | 34 | 32 |
|  | GI-37 | *Pantoea agglomerans* | Soil | 33 | 33 |
|  | GI-17 | *Bacillus aerius* | Leaf | 33 | 32 |
|  | GI-14 | *Pseudomonas glycinis* | Leaf | 32 | 28 |
|  | GI-25 | *Pantoea cypripedii* | Snow | 31 | 28 |
|  | GI-16 | *Bacillus clarus* | Leaf | 28 | 11 |
|  | GI-51 | *Pseudomonas mandelii* | Snow | 19 | 29 |

We examined the phylogenetic distribution of the AF-degrading taxa that we found in our screens. Interestingly, the majority of isolates we found (21 out of 30) belonged to the phylum Pseudomonadota. The second prevalent taxonomic category was the phylum Firmicutes, due to the number of *Bacillus* species that were identified (5 out of 30). Overall, the distribution of isolates across many phyla indicates a broad ability of bacteria to degrade aflatoxin, without specific taxa-based indications for this ability. Notably, species that showed the strongest AF degradation performance did not group together and were dispersed throughout the phylogenetic tree (Fig. 3, green). However, the one isolate that showed no degradation in starch was taxonomically distant from other isolates (Fig. 3, red).

**Figure 3.**
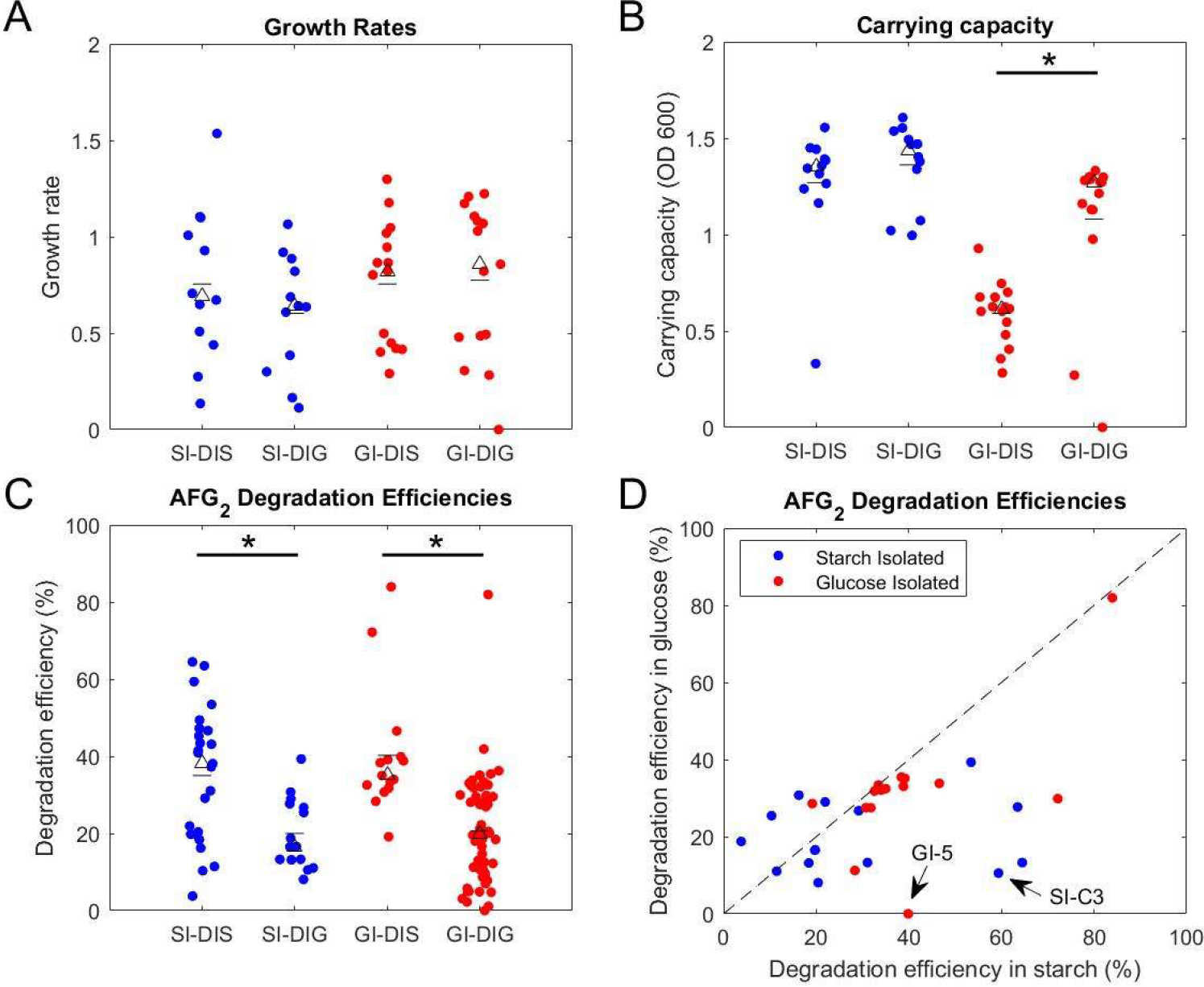
AFG_2_ degradation is improved for glucose-isolated strain when tested in starch medium. Isolates were tested for their growth and AF degradation efficiency when grown in starch and glucose defined media. Starch isolated strains (SI) are shown in blue and glucose isolated strains (GI) are shown in red. Testing in starch medium is indicated by DIS and testing in glucose medium is indicated by DIG. A) Growth rates. B) Carrying capacity. C) Degradation efficiency, shown as percent AF degraded in 48 hours. In A-C, the marked triangles indicate the group’s median, while the marked dash indicates the group’s mean. D) Degradation efficiencies for each isolate in both media. The dotted line represents the same efficiency between the two media. Each dot is the mean of 2 replicates per culturing condition. * = p<0.05, Mann-Whitney U test.

### Starch medium, compared to glucose, improves the degradation performance of isolates

To understand how the environmental carbon source influences degradation, isolates that had undergone 16S rRNA sequencing were tested for aflatoxin degradation performance in the opposite medium from their isolation. Starch isolates had significantly lower degradation efficiency when tested in glucose medium (Fig. 3C, blue), and glucose isolates had significantly increased degradation efficiency when tested in starch (Fig. 3C, red). Additionally, when looking at growth characteristics for isolates between the two medium types, growth rates remained similar for both glucose- and starch-isolates (Fig. 3A). While carrying capacity remained the same for starch isolates in both media, glucose isolates had significantly lower carrying capacity in glucose compared to starch (Fig. 3B). Taken altogether, these dynamics of growth and degradation indicate that lower cell density is not the cause of decreased degradation for the starch isolates in glucose medium, and that for glucose isolates, a lower cell density in starch out-perform the higher cell density of glucose culturing. This increase in performance is likely the result of a metabolism shift when moved to a more complex carbon environment rather than impact on growth since growth rates and carrying capacity remained similar. Looking closer at individual isolates in glucose and starch media, we see the effect that testing in a starch medium has on degradation capacity. As an example, isolate GI-5 showed no degradation when tested in glucose but showed about 40% degradation in starch (Fig. 3D, highlighted). Additionally, isolate SI-C3, a newly identified aflatoxin degrader, decreased its degradation from 60% to 28% when moved into glucose (Fig. 3D, highlighted), which indicates that in a screen using glucose, this new degrader would likely not have been identified. Overall, the fraction of strains that showed higher degradation in starch versus glucose was 73% (11 out of 15) among starch isolates and 93% (14 out of 15) among glucose isolates (Fig 3D).

### Isolates show detoxification ability on other aflatoxins

One key function of a good aflatoxin degrader is the ability to degrade different aflatoxin types. Previous data focused on AFG_2_ since its stronger fluorescence is more reliably detected in our degradation assay. Here, we tested both starch and glucose isolates for their AFB_1_ degradation efficiency to understand the relationship between degradation of these two aflatoxin types. For both sets of isolates in starch medium, there is a linear relationship between AFG_2_ and AFB_1_ degradation, with higher efficiency of AFG_2_ degradation (Fig. 4). This trend is less apparent in the glucose medium for starch isolates, with R^2^ = 0.1799 (Fig. 4B). For glucose isolates tested in glucose medium, the relationship between AFG_2_ and AFB_1_ degradation is much stronger, R^2^= 0.7528 (Fig. 4A). Generally, we find that the best degraders of AFB_1_ are the same as the best AFG_2_ degraders (Fig. 4). Additionally, when comparing the overall performance of isolates on AFB_1_ in starch and glucose media, a similar trend to AFG_2_ is seen in that testing in starch significantly increases degradation efficiency compared to glucose (Fig. S1). The ability of these isolates to degrade both types of aflatoxin in a linear association confirms that the use of AFG_2_ in our assays and screens is adequately representative of AFB_1_ degradation performance.

**Figure 4.**
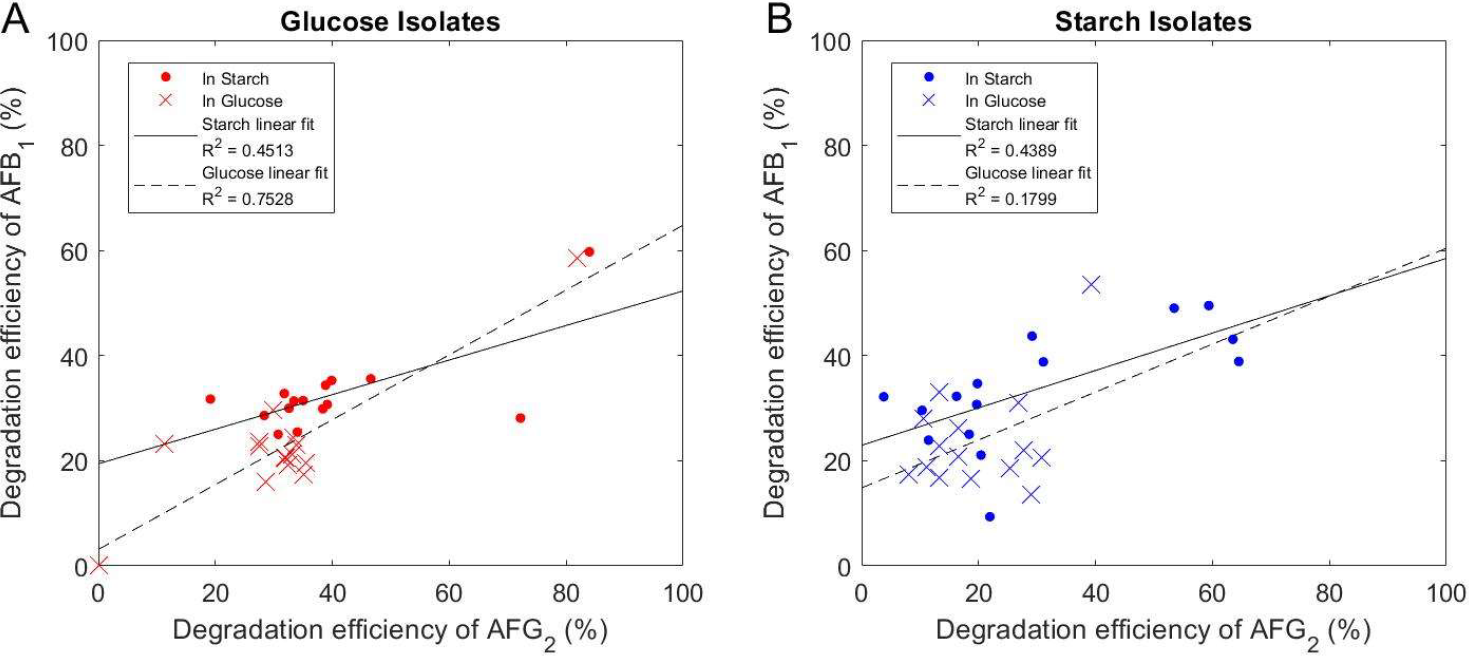
AFB1 degradation by isolates correlates linearly to AFG_2_ degradation efficiency. Isolates were tested for their degradation efficiency on two types of aflatoxin, AFB_1_ and AFG_2_, when grown in starch and glucose defined media. Degradation efficiency is shown as percent AF degraded in 48 hours for A) glucose isolated strains and B) starch isolated strains. Dots (•) represent testing in starch medium and crosses (×) represent testing in glucose medium. Each point is the mean of 2 replicates per culturing condition.

## Discussion

We investigated the possibility of using starch (instead of glucose) as the main carbon source in the growth medium to identify aflatoxin degraders from environmental samples. In this process, we identified new degrader species and found that starch in the environment resulted in an improved degradation phenotype for most isolates. Degradation levels varied in each isolate; however, generally, starch led to higher degradation levels compared to glucose. Additionally, the starch screen allowed for a more streamlined identification of AF degrader species, where growth on starch as the sole carbon source primed candidates for degradation of aflatoxin and facilitated the screening for better degraders.

Of importance in the data shown is how environmental carbon source can change the degradation profiles of certain species. The improvement to degradation potentials by isolates when only moved into a more complex carbon environment implicates a possible regulatory change and/or metabolic shift in the microbes that helps facilitate AF degradation. Using a more complex carbon source as a screen for AF degradation is an intuitive approach. By supplying the cells with a complex carbon as its sole carbon source, we are steering strains that can switch/adapt their metabolism toward complex carbons, and in the process, those that can break down AF. Further studies into the mechanisms behind this process are needed to control and improve this function for practical implementation.

Tested isolates were collected from areas that were not at predisposed risk for AF contamination and they likely did not have prior exposure to AF in the natural environment. The ability of these isolates to degrade AF indicates that the degradation capacity is not necessarily rare among bacterial species. Additionally, the taxonomic breakdown of the analyzed isolates shows a fairly diverse array of species that possess AF degradation ability, further indicating that AF degradation ability can arise in many species of bacteria.

To identify new species of AF degraders, our findings indicate using growth on starch medium is a good screening method due to its low cost, higher percentage of good degrader strains, and better outcome of strains with broader environmental working conditions. Downstream, utilizing a starch screen for samples that have a higher probability of pre-exposure to aflatoxins will be beneficial in finding new degrader strains that possess a high degradation ability. In addition to identifying new degrader strains, it will be important for the application of this degradation capacity to further explore the underlying mechanisms of degradation, such as metabolic pathways, enzyme identification, and degradation by-product analysis.

## Methods

### Environmental isolates and culture mediums

Environmental samples were collected from in and around the Chestnut Hill area in Massachusetts. These samples include soil, snow, leaf, tree trunk, doorknob, and phone screen swabs. Sterile DI water was added to the soil to create a suspension. All samples were struck out on standard LB agar and incubated for 1-3 days at room temperature and 28°C. Individual colonies were then inoculated in either glucose or starch defined medium (medium screens performed from separate environmental samples) in a 96-well plate. Isolates were tested for growth via absorbance at OD600 on a BioTek Synergy Mx microplate reader.

Isolates were cultured in defined medium comprised of 1.5 g/L KH_2_PO_4_, 3.8 g/L K_2_HPO_4_ (x 3H_2_O), 1.3 g/L (NH_4_)_2_SO_4_, 3.0g/L sodium citrate (x 2H_2_O), 20.9 g/L MOPS, 1.1 mg/L FeSO_4_, 1 mL/L mixed vitamin solution (2 mg/L of biotin, 2 mg/L of folic acid, 10 mg/L of pyridoxine-HCl, 5 mg/L of thiamine-HCl ×2H2O, 5 mg/L of riboflavin, 5 mg/L of nicotinic acid, 5 mg/L of D-Ca-pantothenate, 0.1 mg/L of vitamin B12, 5 mg/L of p-aminobenzoic acid, and 5 mg/L of lipoic acid), 1 mL/L SL-10 trace elements solution (10 mL/L of HCl (25%; 7.7 M), 1.5 g/L of FeCl2 ×4H2O, 70 mg/L of ZnCl2, 0.1 g/L of MnCl2 ×4H2O, 6 mg/L of H3BO3, 0.19 g/L of CoCl2 ×6H2O, 2 mg/L of CuCl2 ×2H2O, 24 mg/L of NiCl2 ×6H2O, and 36 mg/L of Na2MoO4 ×2H2O), 1 M MgCl_2_ (5 mL), 1 M CaCl_2_ (1 mL), and 10 mL/L mixed amino acid stock (1.6 g/L of alanine, 1 g/L of arginine, 0.4 g/L of asparagine, 2 g/L of aspartic acid, 0.05 g/L of cysteine, 6 g/L of glutamic acid, 0.12 g/L of glutamine, 0.8 g/L of glycine, 1 g/L of histidine monohydrochloride monohydrate, 2 g/L of isoleucine, 2.6 g/L of leucine, 2.4 g/L of lysine monohydrochloride, 0.6 g/L of methionine, 2 g/L of phenylalanine, 2 g/L of proline, 1 g/L of serine, 0.7 g/L of threonine, 0.3 g/L of tryptophan, 0.25 g/L of tyrosine, 2 g/L of valine, 2 g/L of adenine hemisulfate salt, and 2 g/L of uracil), with either glucose (4.0 g/L) or starch (4.0 g/L) as the carbon source. AFG_2_ and AFB_1_ (Cayman Chemical) was dissolved in LC-MS grade methanol to the final concentration of 1 mg/mL.

### Aflatoxin degradation assay

Cells or culture filtrates were aliquoted into sterile microcentrifuge tubes and aflatoxin was added according to desired final concentration (15-30 μg/mL) per well. Samples were arrayed in black glass-bottom 96-well plates (Nunc™ #165305 96-Well Optical Bottom) at a final volume of 150 μL per well. Standard controls of toxin alone (AFG_2_ in fresh medium) and no toxin (cells or filtrate alone) were used. A BioTek Synergy Mx multi-mode microplate reader was used to monitor optical density of cells at 600 nm and fluorescence of aflatoxin at an excitation of 380 nm and emission of 440 nm with a gain of 50. Reads were taken at 5 min intervals over 72 hours (unless otherwise noted). Cultures usually started at an initial OD of 0.01 and were continuously shaking between reads. Typically, 2-3 replicates were used per condition. Sterile water was placed at the peripheral wells of the 96-well plate to contain evaporation.

### PCR and 16S rRNA gene sequencing

Isolates underwent colony PCR for 16S rRNA gene amplification. Cells were taken from agar plates, suspended in 10 μL Milli-Q, and lysed at 98°C for 15 min. Universal primers used for amplification and sequencing were: 27F (5’-AGAGTTTGATCCTGGCTCAG-3’) and 1492R (5’-GGTTACCTTGTTACGACTT-3’). PCR product was sent for Sanger sequencing, both forward and reverse, at Eurofins Genomics. Resulting sequences were analyzed for the consensus sequence between the forward and reverse sequences using the sequence alignment function of BLASTn (NCBI) and then by megablast through BLASTn for closest sequence match for strain identification.

### Phylogenetic tree

Based on BLASTn analysis and the closest match for each isolate, we used the GenBank reference sequences for the creation of the phylogenetic tree through multi-sequence alignment. We created the phylogenetic tree file using the Matlab functions `multialign`, `seqpdist`, and `seqlinkage`. The tree file was annotated using Interactive Tree of Life (iTOL) (30).

### Data analysis and statistics

Raw data from the aflatoxin degradation assay was processed using Matlab generated codes to measure growth and degradation characteristics. Background fluorescence (no toxin control) is subtracted from the readings to remove fluorescence from sources other than the toxin. The readouts are also normalized to fluorescence data from no cell controls to remove the effect of fluorescence loss due to bleaching or other causes over time with an additional normalization to account for fluorescence loss due to cell scattering (Fig. S2). To convert the fluorescence readout to the corresponding toxin concentration, we employ a calibration curve based on measurements of a set of known toxin concentrations (31). After normalization, degradation efficiency is calculated as the percentage of toxin removed during the testing period (48 hours).

### Statistical analysis

To compare the odds ratio between good degraders and poor or non-degraders obtained from the starch versus glucose isolation (Fig. 1), we used Fisher’s exact test using the Matlab function `fishertest`. To compare the growth rates, carrying capacities, and AFG_2_ degradation efficiency between starch-versus glucose-isolates (Fig. 3), we used Mann-Whitney U test using the Matlab function `ranksum`.

## Data availability

All data and analysis files presented in this work will be available from the authors upon request.

## Acknowledgements

This work was partially supported by the National Science Foundation (NSF-CBET) under Grant No. 2103545. NS is supported by a NIFA-AFRI Predoctoral Fellowship from the USDA (Award No. 2021-67034-35108).

## Supplemental figures

**Figure S1.**
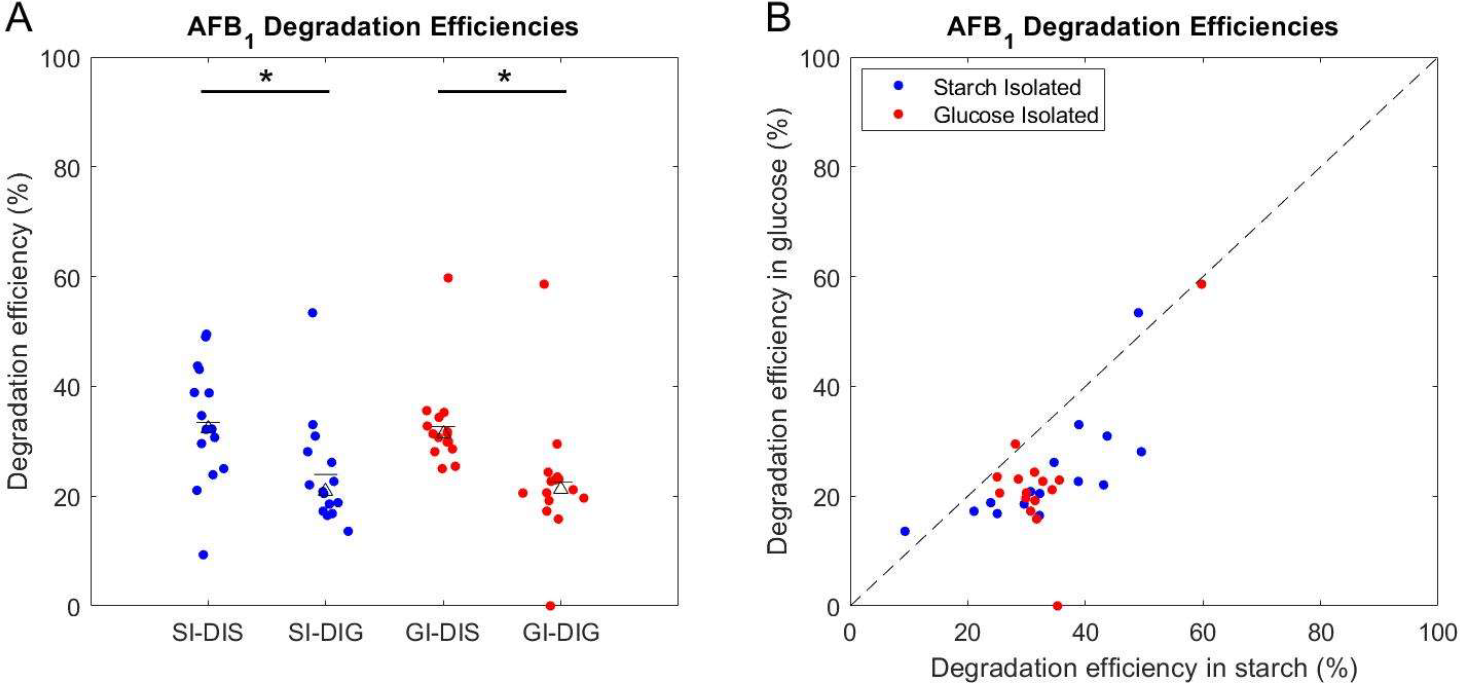
AFB_1_degradation is improved when tested in starch medium. Isolates were tested for their AF degradation efficiency when grown in starch and glucose defined media. Starch isolated strains are shown in blue and glucose isolated strains are shown in red. A) Degradation efficiency, shown as percent AF degraded in 48 hours, grouped by isolation and testing medium. Testing in starch medium is indicated by DIS and testing in glucose medium is indicated by DIG. Degradation efficiency is shown as percent AF degraded in 48 hours. The marked triangles indicate the group’s median, while the marked dash indicates the group’s mean. B) Degradation efficiencies for each isolate in both media. The dotted line represents the same efficiency between the two media. Each dot is the mean of 2 replicates per culturing condition. * = p<0.05, Mann-Whitney U test.

**Figure S2.**
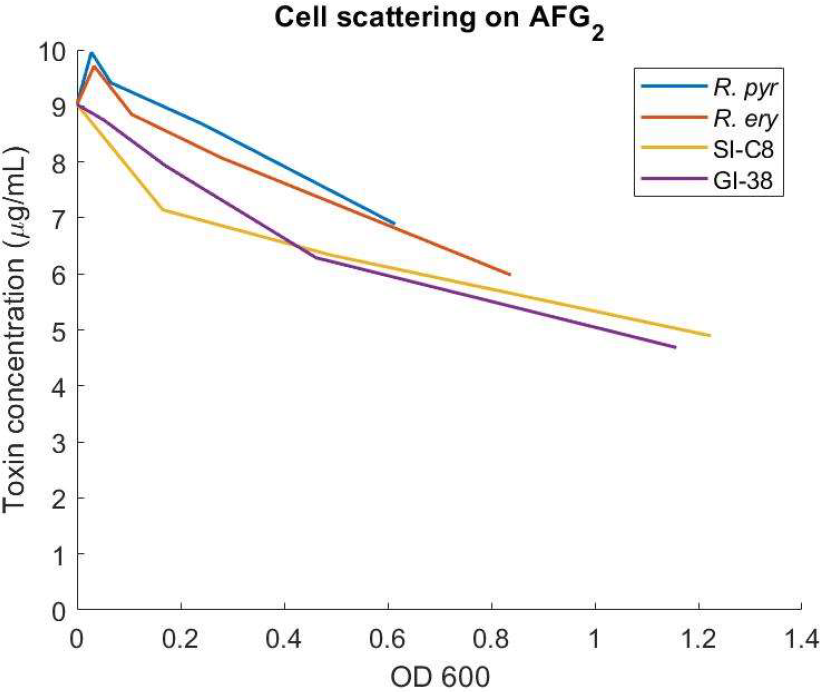
Cell-scattering influences the fluorescence readout of AF. The same concentration of AFG_2_ was added to serial dilutions of cells immediately prior to measuring fluorescence in our FL assay. Readings reflect the effects of fluorescence scattering due to cell density. Linear slope for each representative strain was calculated and averaged to set as the normalization of cell scattering during growth in our FL assay.

